## Supplementary for "Infusing structural assumptions into dimensionality reduction for single-cell RNA sequencing data to identify small gene sets"

*These authors contributed equally.

#### Contents

**S1 Complementary UMAP plots colored by BAE latent dimensions on cortical mouse data** 3

**S2 Cell type correspondence of BAE latent dimensions on cortical mouse data** 4

**S3 Summary of the genes selected per latent dimension on cortical mouse data** 5

**S4 BAE mappings of held out cortical mouse test data** 7

**S5 Comparison of the BAE with supervised componentwise boosting on cortical mouse data** 8

**S6 Gene selection stability analysis on cortical mouse data** 10

**S7 Robustness analysis of structuring latent representations on cortical mouse data** 12

**S8 UMAP embedding of PCA representation of cortical mouse data** 14

**S9 Model comparison for gene selection on cortical mouse data** 15

S10Comparison of different timeBAE encoder initialization strategies 16

S11Complementary UMAP plots for timeBAE analysis of embryoid body data 17

S12timeBAE gene selection comparison 20

S13Simulation of stair-like scRNA-seq-like data 21

S14Simulation study to illustrate how the BAE identifies distinct cell groups and corresponding explanatory genes 21

S15Model comparison of different autoencoder versions on simulated data 24

S16Pseudocodes 32

S17References 34

#### S1 Complementary UMAP plots colored by BAE latent dimensions on cortical mouse data

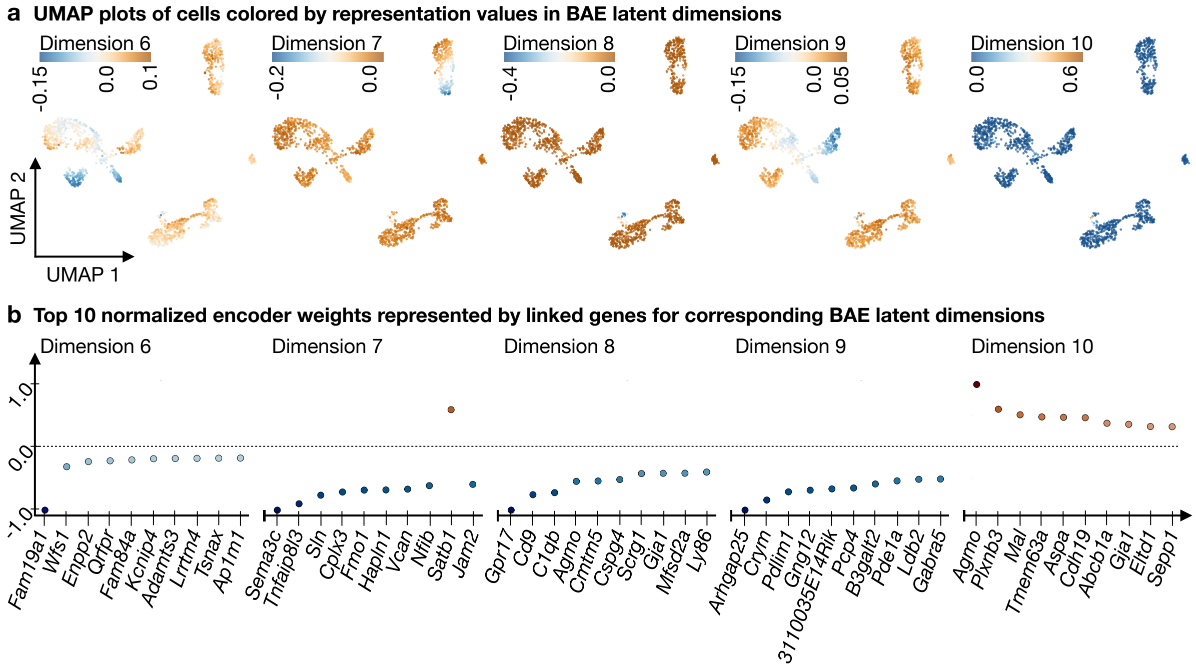


Figure S1: **Additional latent dimensions for the BAE analysis on cortical mouse scRNA-seq data. a** Scatter plots of a two-dimensional UMAP embedding computed using a ten-dimensional BAE latent representation of the cortical neurons. Cells are colored by their representation values in individual BAE latent dimensions (the last five dimensions are shown). **b** Scatter plots show the top ten normalized nonzero BAE encoder weights (represented by linked genes) in decreasing order of absolute value (from left to right) per latent dimension (the last five dimensions are shown). Normalization was performed by dividing each weight by the maximum absolute weight value for the corresponding latent dimension.

#### S2 Cell type correspondence of BAE latent dimensions on cortical mouse data

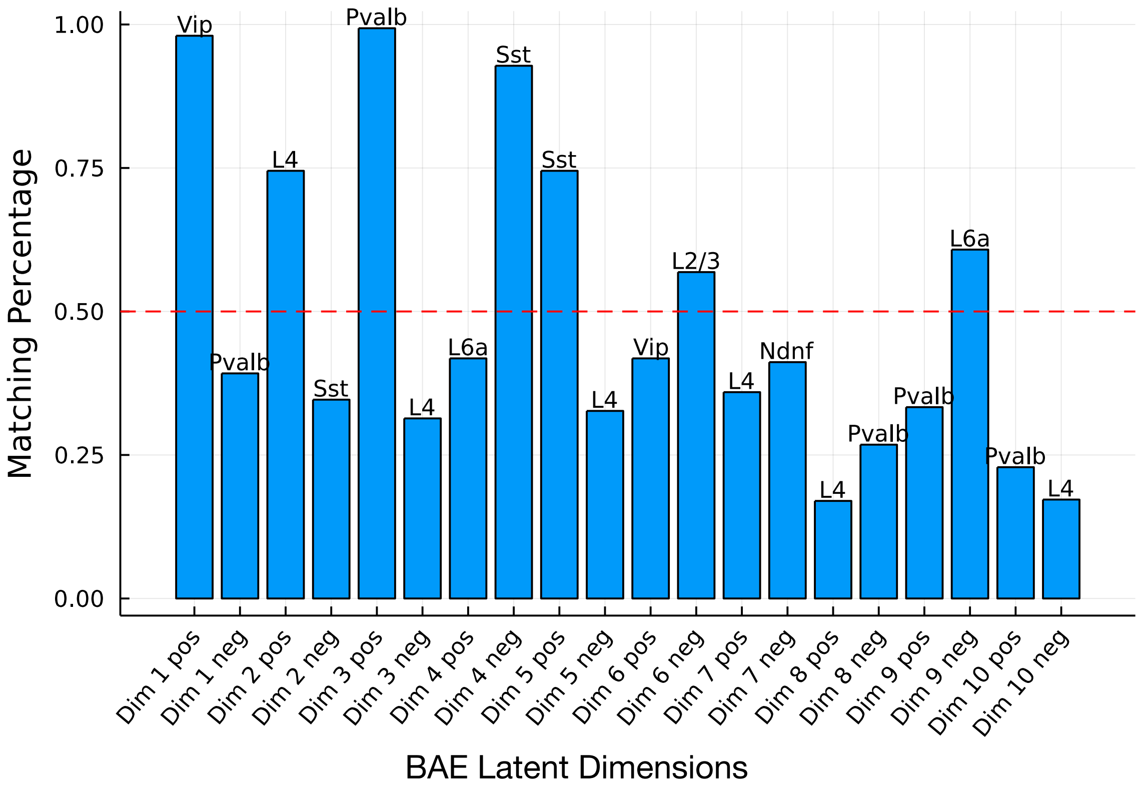


Figure S2: **Comparison of the cell type patterns with the patterns learned in the BAE latent dimensions.** The bar plot shows the percentage of cells belonging to the best matching cell type per latent dimension (see Methods Section 5.8 in the main text). Since cells can have either a positive or a negative representation value in each latent dimension, the positive and negative patterns are compared with the cell type patterns per latent dimension. For example, the most positive cell representations in latent dimension 1 strongly correspond to cells of the type Vip. The red horizontal line indicates the 50% line.

#### S3 Summary of the genes selected per latent dimension on cortical mouse data

During training and starting from a zero initialization, the BAE selects weights in the encoder weight matrix to be updated. Nonzero weights represent a connection between genes and latent dimensions. After training the encoder weight matrix of a BAE is sparse, i.e. there are only a few connections from genes to each latent dimension. In the analysis of the cortical mouse data, the encoder weight matrix consisted of 10 latent dimensions and 1500 variables, i.e., genes. Out of the 15000 weights, 495 weights were nonzero (3.3%). The following tables provide information about the distribution of nonzero weights across the latent dimensions (Table 1) and the top selected genes, i.e., change-point genes per latent dimension (Table 2). Furthermore, the changes of the BAE encoder weight values can be tracked across training epochs. The dynamics of the encoder weights (coefficients) are presented in Figure S3.

| Latent dimension | Number of BAE selected genes (nonzero weights) | Percentage of selected genes (nonzero weights) | Number of change-point genes (nonzero weights) | Percentage of change-point genes (nonzero weights) |
|---|---|---|---|---|
| **1** | 47 | 3.13 | 1 | 0.07 |
| **2** | 54 | 3.6 | 2 | 0.13 |
| **3** | 53 | 3.53 | 1 | 0.07 |
| **4** | 57 | 3.8 | 1 | 0.07 |
| **5** | 56 | 3.73 | 1 | 0.07 |
| **6** | 58 | 3.87 | 1 | 0.07 |
| **7** | 47 | 3.13 | 2 | 0.13 |
| **8** | 36 | 2.4 | 1 | 0.07 |
| **9** | 57 | 3.8 | 1 | 0.07 |
| **10** | 30 | 2.0 | 1 | 0.07 |

Table 1: **Summary of the number of genes selected per latent dimension. First column:** Number of genes selected out of the total number of genes (1500) during training per latent dimension. **Second column:** Percentage of genes selected. **Third column:** Number of change-point genes (see Methods Section 5.9 in the main text) indicating the top selected genes w.r.t. the absolute values of the corresponding encoder weights per latent dimension. **Fourth column:** Percentage of change-point genes.

| Latent dimension | Changepoint gene 1 | Changepoint gene 2 |
|---|---|---|
| **1** | *Vip* | |
| **2** | *Rorb* | *Pamr1* |
| **3** | *Pvalb* | |
| **4** | *Galntl6* | |
| **5** | *Chodl* | |
| **6** | *Fam19a1* | |
| **7** | *Sema3c* | *Tnfaip8l3* |
| **8** | *Gpr17* | |
| **9** | *Arhgap25* | |
| **10** | *Agmo* | |

Table 2: **Change-point genes per latent dimension.** The table shows the change-point genes per latent dimension (see Methods section 5.9 in the main text), sorted by the corresponding absolute values of the encoder weights. The maximum number of change-point genes per latent dimension in the mouse cortical data is 2 and the patterns observed in the latent dimensions are predominantly driven by the change-point genes.

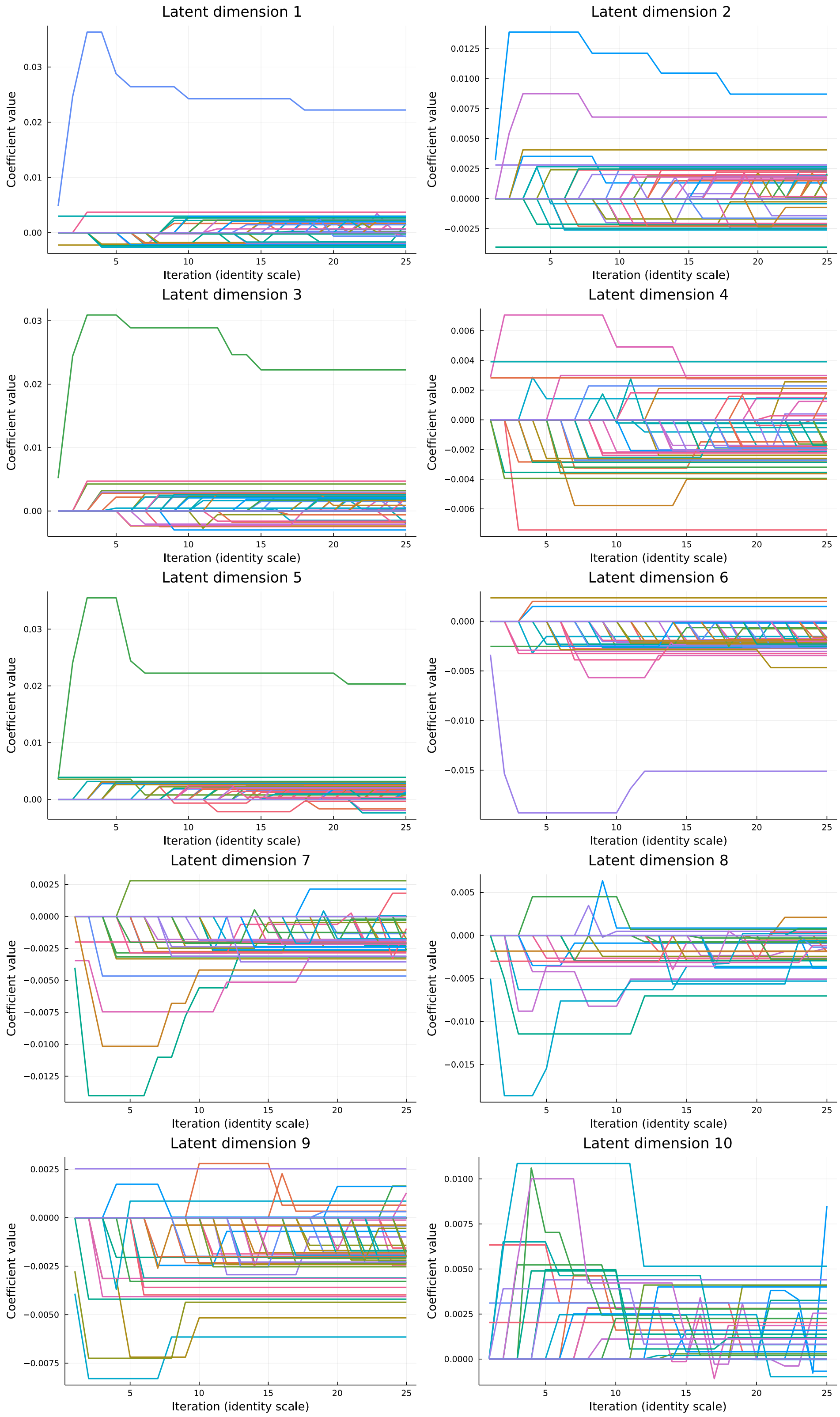


Figure S3: **Dynamics of BAE encoder weights across training epochs.** The plots show the evolution of individual BAE encoder weights (coefficients) across training epochs per latent dimension. Each line represents a specific weight corresponding to a gene.

#### S4 BAE mappings of held out cortical mouse test data

We mapped the held-out test data (200 randomly selected cells from the 1,525 cortical mouse cells) to the latent space using the trained encoder of the BAE. To assess whether the test cells were accurately mapped to regions corresponding to their cell types in the training data, we predicted the test cell types by identifying the 10 closest cells in the latent space from the training data (based on Euclidean distance) and assigning the most frequent cell type among them (see Methods Section 5.10 in the main text). This approach yielded a prediction accuracy of 94% when compared to the true labels of the test data. Figure S4 illustrates two-dimensional UMAP embeddings of the BAE training and test data representations.

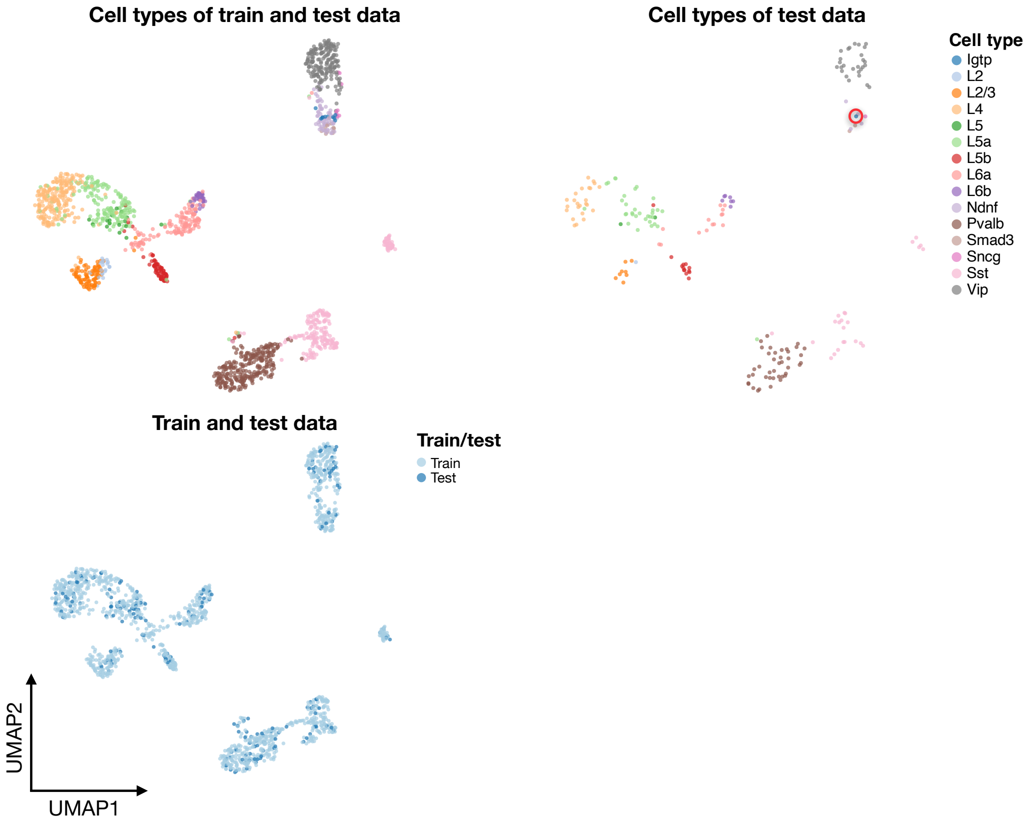


Figure S4: **UMAP plots cortical mouse data (train and test data). Top left:** Scatter plot of a two-dimensional UMAP embedding of the BAE latent representation of the neurons colored by cell types (train and test data). **Top right:** Scatter plot of a two-dimensional UMAP embedding of the BAE latent representation of the neurons in the test data colored by cell types. An Igtp cell was added to the test data to ensure a consistent color coding. This cell is highlighted with a red circle. **Bottom left:** Scatter plot of a two-dimensional UMAP embedding of the BAE latent representation of the neurons in the test data, colored by training and test IDs.

#### S5 Comparison of the BAE with supervised componentwise boosting on cortical mouse data

We compared the learned representation of the BAE with that of the supervised comp$L_2$Boost algorithm, which is at the core of the optimization of the BAE. For applying the comp$L_2$Boost algorithm to the cortical mouse data, we use the cell type annotations from [1] and created a one-hot encoded version of the cell labels that were stored in a matrix where the rows correspond to the cells and the columns to the different cell types. Then, for each cell type in the data we applied comp$L_2$Boost for respectively 100 boosting steps with a step size of 0.02 using the standardized training data and the corresponding column of the one-hot matrix.

Figure S5 shows a scatter plot of a two-dimensional UMAP embedding of the neurons using the comp$L_2$Boost representation analogous to Figure 2. In each plot, the embedding is colored according to the model prediction for one specific cell type, i.e., where the corresponding column in the one-hot encoded cell type matrix was used as the responses.

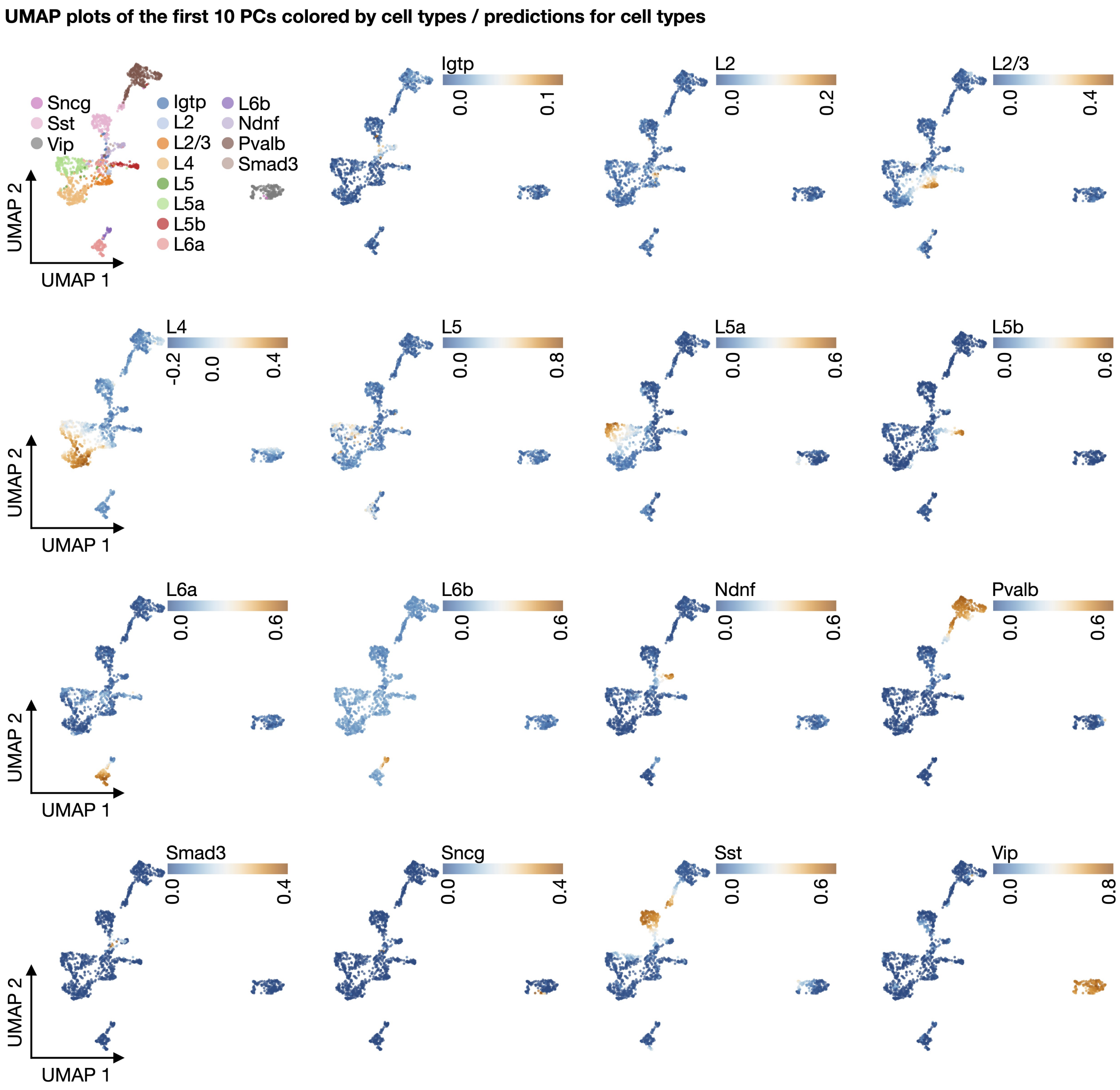


Figure S5: **UMAP plots of cortical mouse data colored by comp$L_2$Boost predictions. Top left:** Scatter plot of a two-dimensional UMAP embedding of the neurons colored by cell types. The UMAP embedding was computed using the concatenated predictions of a componentwise boosting model applied sequentially to each column of a one-hot encoding of the cell labels. **Other plots:** Scatter plots show the same UMAP embedding colored by the predictions of the componentwise boosting model for each cell type. The plots are labeled by the cell type used to obtain the predictions, respectively.

#### S6 Gene selection stability analysis on cortical mouse data

We analyzed the stability of gene selection in the BAE model using the cortical mouse data, as detailed in Methods Section 5.11 of the main text. The results are presented in a bar plot (Figure S6). Additionally, we examined genes selected in 80% or more of the 30 runs (Table 3). Across the 30 training runs, the BAE encoder weight matrices had an average of 96.25% zero elements, with a 95% confidence interval of [96.21%, 96.29%], rounded to two decimal places.

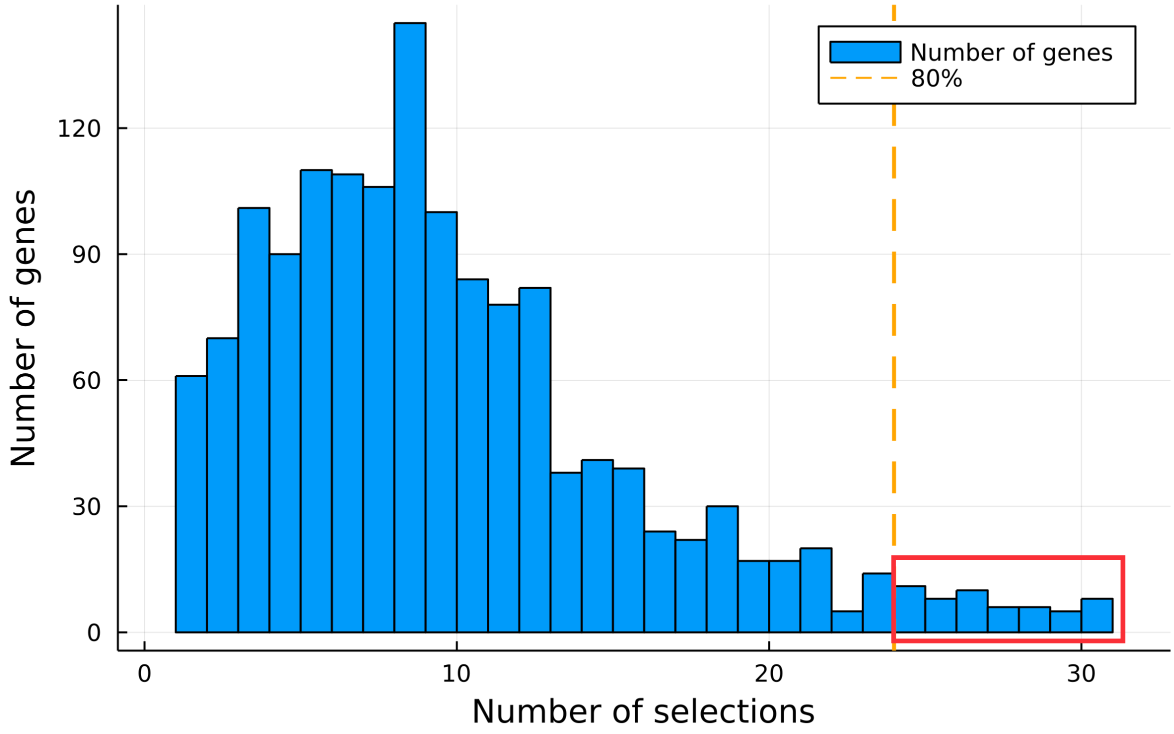


Figure S6: **Distribution of BAE gene selections.** Bar plot represents the number of individual genes (y-axis) with respect to the number of selections across different BAE training runs with different decoder parameter initializations (x-axis). The total number for which a gene could be selected was 30. The orange dashed vertical line reflects the 80% threshold of the number of selections and the red box marks the number of selections that lie above this threshold, reflecting genes that were selected in at least 24 of the 30 runs (see Table 3).

| | Gene | Number of selections | Percentage of selections | Gene type |
|---|---|---|---|---|
| **1** | *Sst* | 30 | 100 | Neural marker |
| **2** | *Cplx3* | 30 | 100 | Neural marker |
| **3** | *Fmo1* | 30 | 100 | |
| **4** | *Slco1a4* | 30 | 100 | Nonneural marker |
| **5** | *Cyba* | 30 | 100 | |
| **6** | *Chodl* | 30 | 100 | Neural marker |
| **7** | *Dgkb* | 30 | 100 | Neural marker |
| **8** | *Agmo* | 30 | 100 | |
| **9** | *Tnfaip8l3* | 29 | 96.66 | Neural marker |
| **10** | *Lrrtm4* | 29 | 96.66 | |
| **11** | *Pdlim1* | 29 | 96.66 | |
| **12** | *Cd9* | 29 | 96.66 | Nonneural marker |
| **13** | *A930038C07Rik (Ndnf)* | 29 | 96.66 | Neural marker |
| **14** | *Pvalb* | 28 | 93.33 | Neural marker |
| **15** | *Vtn* | 28 | 93.33 | Nonneural marker |
| **16** | *Fam84a* | 28 | 93.33 | |
| **17** | *Reln* | 28 | 93.33 | Neural marker |
| **18** | *Plxnb3* | 28 | 93.33 | |
| **19** | *Calb1* | 28 | 93.33 | Neural marker |
| **20** | *2310042E22Rik (Teddm3)* | 27 | 90 | Neural marker |
| **21** | *Myh4* | 27 | 90 | Neural marker |
| **22** | *Cspg4* | 27 | 90 | Nonneural marker |
| **23** | *Rorb* | 27 | 90 | Neural marker |
| **24** | *Olfml1* | 27 | 90 | |
| **25** | *Gpr37l1* | 27 | 90 | |
| **26** | *Anax3* | 26 | 86.66 | |
| **27** | *Ly86* | 26 | 86.66 | Nonneural marker |
| **28** | *C1qb* | 26 | 86.66 | Nonneural marker |
| **29** | *Gpr17* | 26 | 86.66 | Nonneural marker |
| **30** | *Gng12* | 26 | 86.66 | |
| **31** | *Vip* | 26 | 86.66 | Neural marker |
| **32** | *Mas1* | 26 | 86.66 | |
| **33** | *Foxp2* | 26 | 86.66 | Neural marker |
| **34** | *Eltd1* | 26 | 86.66 | Nonneural marker |
| **35** | *Wfs1* | 26 | 86.66 | Neural marker |
| **36** | *Enpp2* | 25 | 83.33 | Neural marker |
| **37** | *Nfia* | 25 | 83.33 | |
| **38** | *Mag* | 25 | 83.33 | Nonneural marker |
| **39** | *Myh13* | 25 | 83.33 | Neural marker |
| **40** | *Myh8* | 25 | 83.33 | Neural marker |
| **41** | *Cpne6* | 25 | 83.33 | |
| **42** | *Gng11* | 25 | 83.33 | |
| **43** | *Slc38a3* | 25 | 83.33 | |
| **44** | *Gabra1* | 24 | 80 | Neurotransmitter receptor |
| **45** | *Gja1* | 24 | 80 | Nonneural marker |
| **46** | *Ecscr* | 24 | 80 | |
| **47** | *Slco1c1* | 24 | 80 | Nonneural marker |
| **48** | *6330527O06Rik (Lamp5)* | 24 | 80 | Neural marker |
| **49** | *Olig2* | 24 | 80 | |
| **50** | *Pcp4* | 24 | 80 | |
| **51** | *Spock1* | 24 | 80 | |
| **52** | *Cmtm5* | 24 | 80 | |
| **53** | *Arhgap25* | 24 | 80 | Neural marker |
| **54** | *Jam2* | 24 | 80 | |

Table 3: **Most frequently selected genes.** The table lists genes selected by the BAE in at least 80% of the 30 runs. Columns display the gene names, number of selections, percentage of selections, and their classification based on [1]. Genes without a classification were not listed in [1].

### S7 Robustness analysis of structuring latent representations on cortical mouse data

Details of the robustness analysis can be found in Methods Section 5.12 of the main text, while a summary of the structured gene replacement procedure is provided in Table 4.

The learned structure of the BAE latent representation was assessed by visually inspecting 2D UMAP projections of cell representations to gain insights into the latent patterns (Figure S7). At a gene replacement rate of 2.87% (removal of the top 43 genes identified in the gene selection stability analysis, Table 3), a significant portion of neural markers was already missing. At 13.53%, the BAE struggled to separate cells from different cortical layers into distinct clusters while still preserving the structure of GABAergic neurons. At 61.07%, layer-specific separation further deteriorated, though GABAergic neuron structure was roughly maintained. At the final replacement rate of 97.13%, the model barely distinguished GABAergic from glutamatergic neurons, yet interestingly, *Chodl*-expressing neurons (a subset of *Sst*-expressing neurons) often formed distinct clusters, even in the absence of the *Chodl* marker.

Overall, the BAE demonstrated a robust preservation of the core neural cell structure, even when a substantial fraction of informative genes was replaced with noise.

| Selection frequency threshold | 0% | 10% | 20% | 30% | 40% | 50% | 60% | 70% | 80% | 90% | 100% |
|---|---|---|---|---|---|---|---|---|---|---|---|
| Number of genes replaced | 1457 | 1225 | 916 | 565 | 321 | 203 | 127 | 73 | 43 | 19 | 0 |
| Percentage of genes replaced | 97.13% | 81.67% | 61.07% | 37.67% | 21.4% | 13.53% | 8.74% | 4.87% | 2.87% | 1.27% | 0.0% |

Table 4: **Systematic replacement of informative genes with noise genes in cortical mouse data.** The table below summarizes the number of highly variable genes (out of 1500) in the cortical mouse dataset that were replaced with noise genes. The replacement was based on the selection frequency threshold, which represents the percentage of runs in which a gene was selected during the gene selection stability analysis (see Supplementary Section S6 and Supplementary Table 3). Genes with selection frequencies above the threshold were replaced. In total, 1457 of the 1500 highly variable genes were selected by the BAE across all runs.

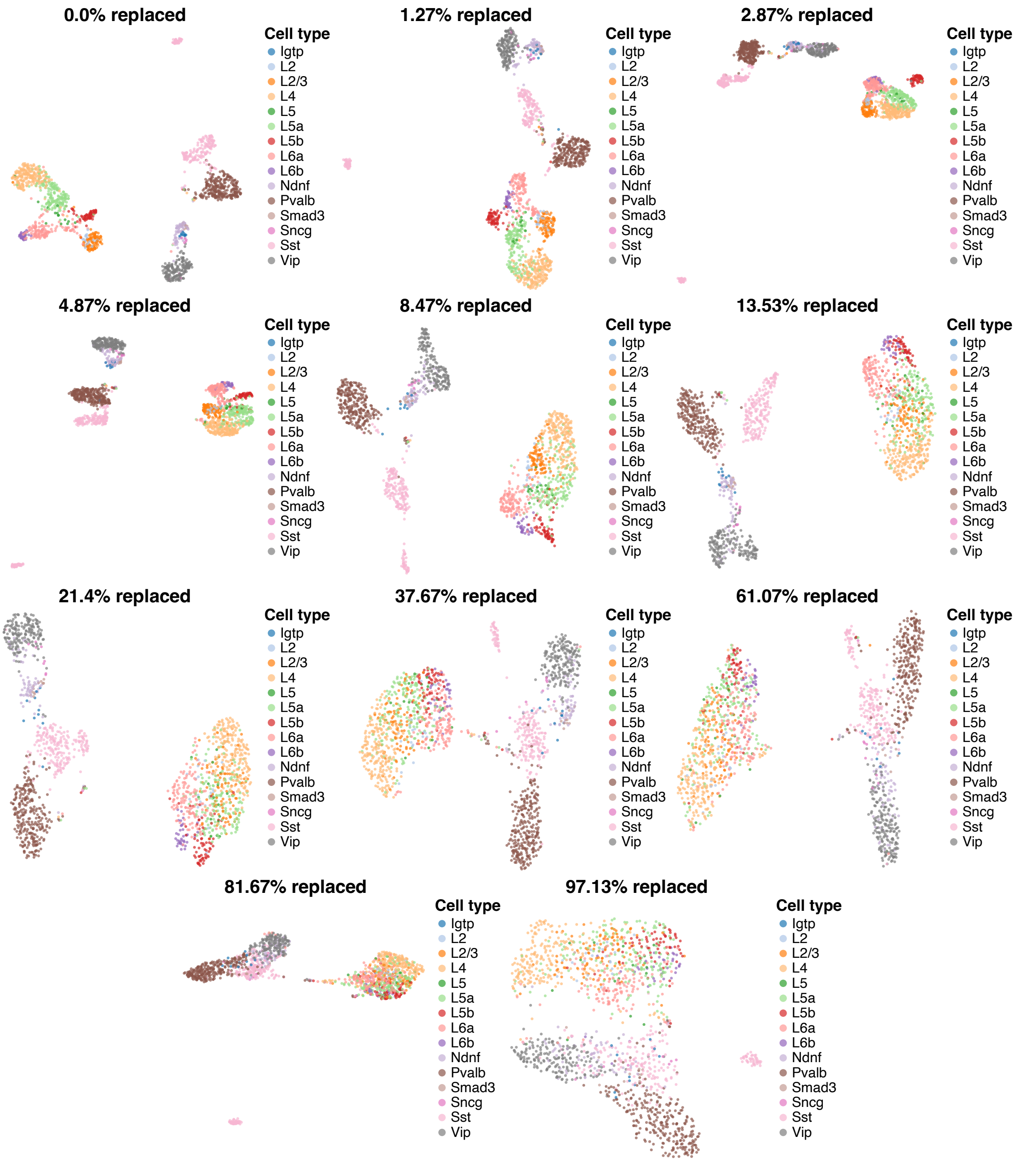


Figure S7: **BAE robustness analysis of structuring latent representations.** Scatter plots of 2D UMAP embeddings of BAE latent representations for the 1525 neural cells from the cortical mouse data are shown at varying levels of informative gene replacement with noise genes. The original dataset consists of 1500 highly variable genes (for details, see Methods Section 5.19 in the main text). All datasets consistently included 1,500 genes.

#### S8 UMAP embedding of PCA representation of cortical mouse data

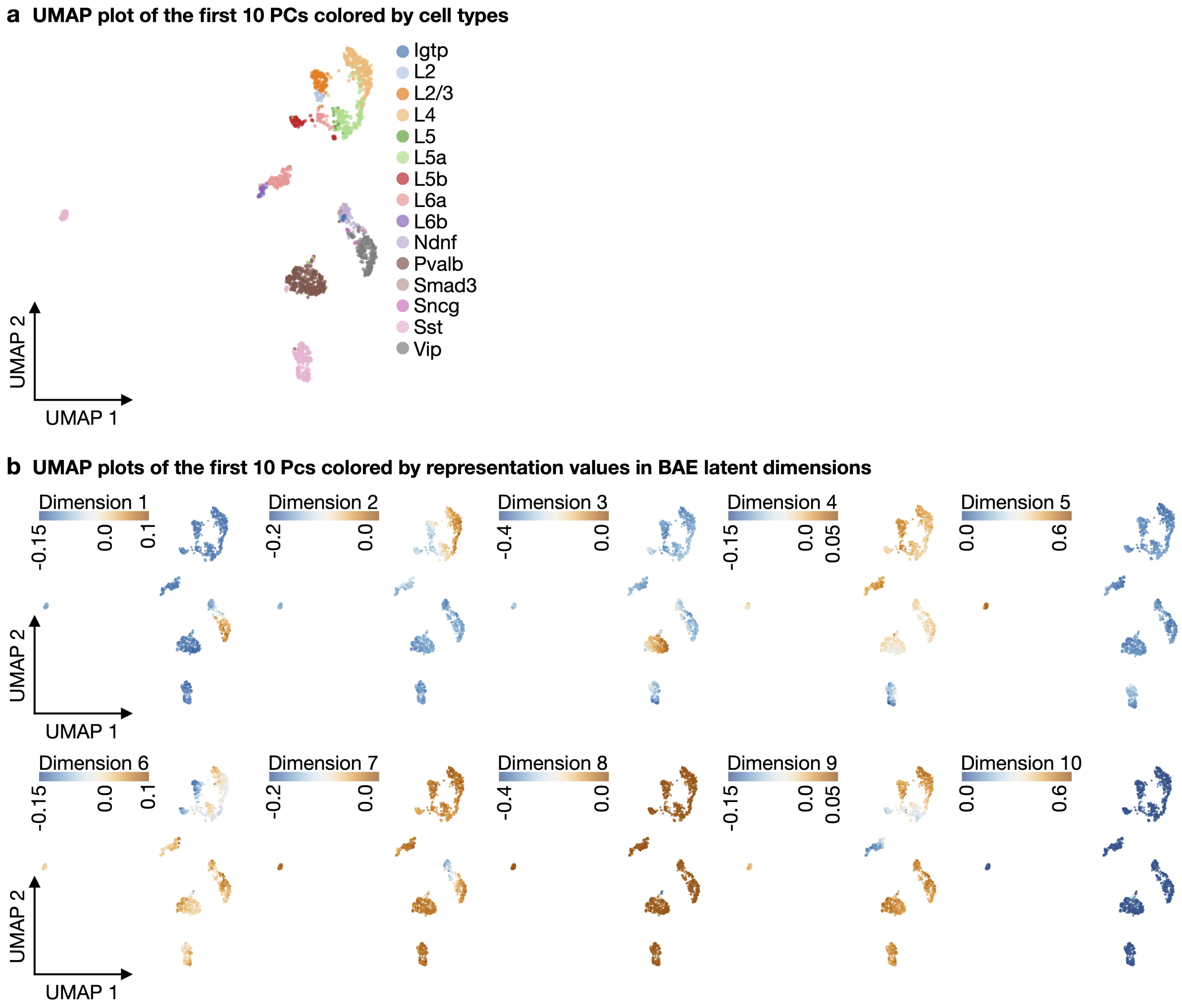


Figure S8: **Colored UMAP embeddings of the PCA latent representation of adult mouse neurons. a** Scatter plot of a two-dimensional UMAP embedding of the first ten PCs of the neurons colored by cell types. **b** Scatter plots of the same UMAP embedding colored by the learned representation values of cells in the ten latent dimensions of the BAE.

#### S9 Model comparison for gene selection on cortical mouse data

The BAE was compared to sparsePCA [2] and scPNMF [3] in terms of gene selection, specifically the sparsity of the loading matrices and the BAE encoder weight matrix. Consistent with the gene selection stability analysis (see Supplementary Section S6), we trained the BAE 30 times under identical training conditions but with different initializations of the decoder parameters. Similarly, sparsePCA and scPNMF were each run 30 times with varying seed parameters. The results, including sparsity levels (measured as the number of zero elements in the BAE encoder weight matrices and loading matrices), are summarized in Table 5. For a detailed explanation of the analysis, refer to Methods Section 5.14 in the main text.

Under the given conditions, the BAE produced encoder weight matrices with the highest sparsity level, achieving a mean percentage of zeros at 96.25%. On average, 448.57 genes were selected per run, representing 29.90% of the 1,500 genes. In contrast, sparsePCA and scPNMF exhibited (close to) zero variability across individual runs. The loading matrices for sparsePCA and scPNMF had an average of approximately 41% and 45% zero elements, respectively, resulting in significantly more nonzero elements compared to the BAE encoder weight matrices. Consequently, almost all genes were connected to at least one component by a nonzero element in the loading matrices, with sparsePCA leaving only one gene unconnected. The average number of selected genes per run was 1,499 for sparsePCA and 1,500 for scPNMF.

Note, that the sparsity level of the different methods is controlled by model-specific hyperparameters for sparsePCA and scPNMF [2, 3]. For comparison, we used the default hyperparameter settings for each approach. Additionally, factors such as the number of training epochs or the step size of the boosting component for optimizing the encoder parameters can influence the sparsity level of the resulting BAE encoder weight matrix, as discussed in detail in Methods Section 5.3 and 5.4 in the main text.

| Model | Mean percentage of zeros (loading matrix/ encoder weight matrix) | 95% CI of percentage of zeros | Mean number of selected genes | Mean percentage of selected genes | Number of genes selected in all runs |
|---|---|---|---|---|---|
| **BAE** | 96.25% | [96.21%, 96.29%] | 448.57 | 29.90% | 1457 |
| **sparsePCA** | 40.85% | – | 1499 | 99.93% | 1499 |
| **scPNMF** | 44.76% | – | 1500 | 100% | 1500 |

Table 5: **Gene selection comparison of the BAE with sparsePCA and scPNMF.** The table summarizes the percentages of zero elements in the BAE encoder weight matrix and the loading matrices for sparsePCA and scPNMF across 30 training runs. Additionally, it reports the number of selected genes, where a gene was considered selected in a run if it had at least one nonzero element across the components in the corresponding BAE encoder weight matrix or loading matrices. 95% confidence intervals for sparsePCA and scPNMF are omitted because there was no variability in the nonzero elements across runs.

#### S10 Comparison of different timeBAE encoder initialization strategies

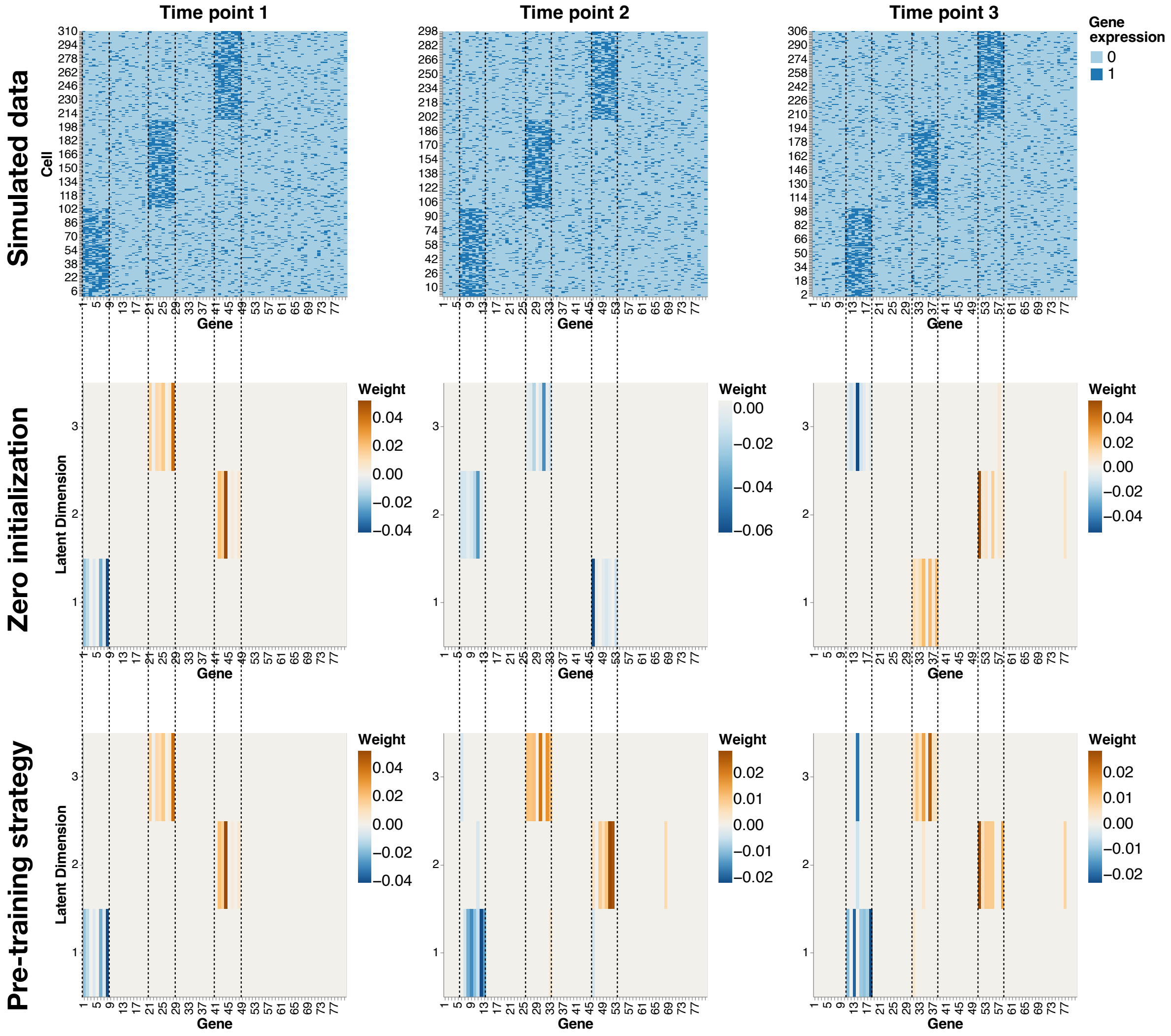


Figure S9: **timeBAE results on simulated data using different encoder initialization strategies. Top row:** Simulated binary count data representing three different cell groups evolving over three different time points. **Middle row:** BAE encoder weight matrices learned at each time point if all encoder weights were set to zero at the beginning of training for each BAE model. Although cell group patterns are captured in disentangled latent dimensions at each time point, no linkage of latent dimensions across time is achieved. **Bottom row:** The timeBAE encoder weight matrices are learned at each time point when the pre-training strategy is used. In this case, the model is able to capture cell group patterns in disentangled latent dimensions, while also linking corresponding latent dimensions across time.

#### S11 Complementary UMAP plots for timeBAE analysis of embryoid body data

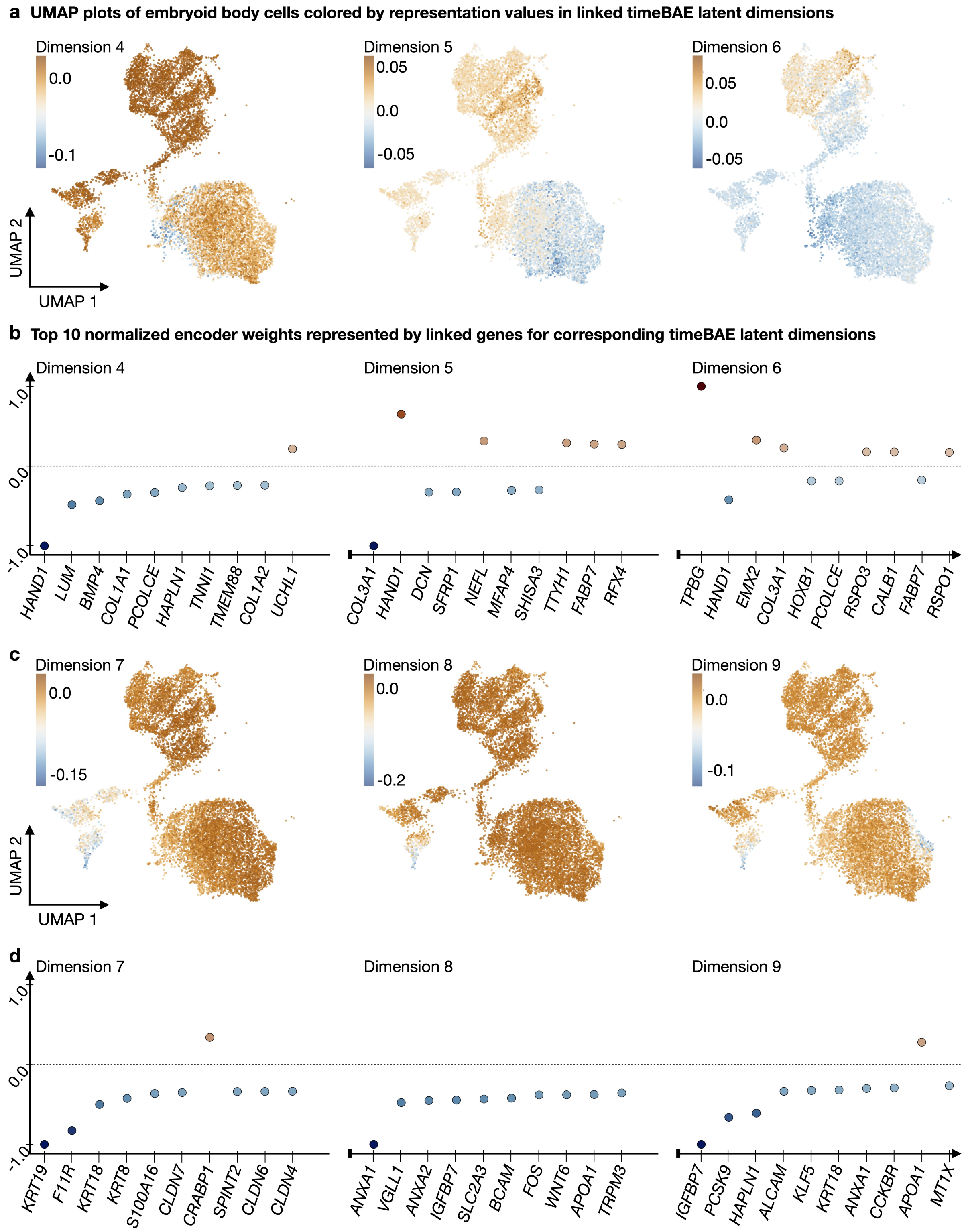


Figure S10: **Additional latent dimensions (4-9) for the timeBAE approach on embryoid body data.a** Scatter plots of a 2D UMAP embedding of the PCA representation of the trimmed embryoid body data. Plots are colored by the representation values of cells in linked timeBAE latent dimensions across time (dimensions 4-6 are shown). **b** Scatter plots show the top ten normalized nonzero timeBAE encoder weights (represented by linked genes) in decreasing order of absolute value (from left to right) per latent dimension (dimensions 4-6 are shown). Normalization was performed by dividing each weight by the maximum absolute weight value for the corresponding latent dimension. The first dimension has only seven corresponding nonzero weights. **c** Colored scatter plots of the UMAP embedding (dimensions 7-9 are shown). **d** Scatter plots of normalized coefficients (dimensions 7-9 are shown)

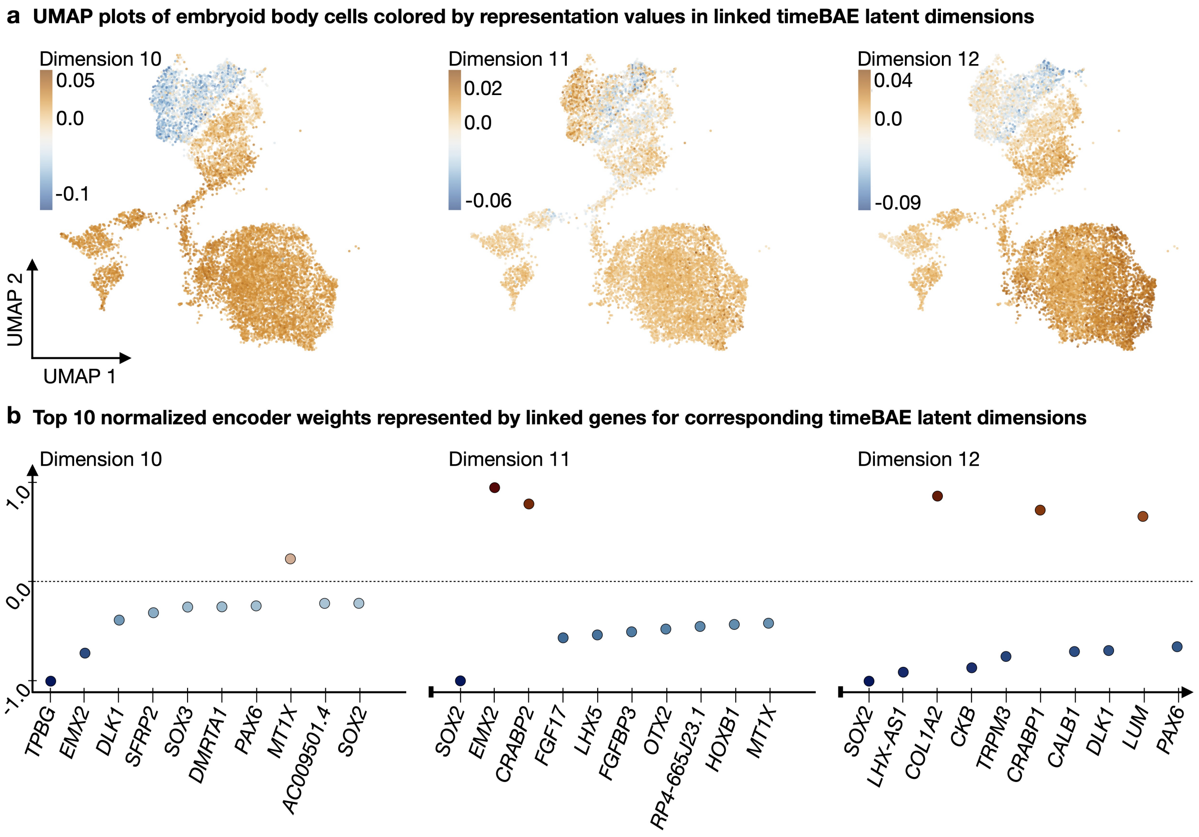


Figure S11: **Additional latent dimensions (10-12) for the timeBAE approach on embryoid body data.a** Scatter plots of a 2D UMAP embedding of the PCA representation of the trimmed embryoid body data. Plots are colored by the representation values of cells in linked timeBAE latent dimensions across time (dimensions 10-12 are shown). **b** Scatter plots show the top ten normalized nonzero timeBAE encoder weights (represented by linked genes) in decreasing order of absolute value (from left to right) per latent dimension (dimensions 10-12 are shown). Normalization was performed by dividing each weight by the maximum absolute weight value for the corresponding latent dimension. The first dimension has only seven corresponding nonzero weights.

#### S12 timeBAE gene selection comparison

In a setting, where both time point and cluster annotations of cells are available, the timeBAE results can be compared to a linear regression analysis. The time point and cluster annotations can be used to determine the most significant genes and compare them to the top genes selected by the timeBAE. The supervised linear modeling approach and comparison process are described in detail in Methods Section 5.16 of the main text.

| C = Cluster<br>T = Time Point | Visually matching timeBAE latent dimension | Number of selected genes timeBAE | Number of selected genes linear model | Number of gene overlap | Percentage of gene overlap |
|---|---|---|---|---|---|
| **All** | – | 87 | 124 | 59 | 67.82% |
| **C1 T1** | 4 | 10 | 36 | 5 | 50.0% |
| **C1 T2** | 5 | 12 | 56 | 7 | 58.33% |
| **C1 T3** | 6 | 11 | 39 | 5 | 45.45% |
| **C2 T1** | 10 | 12 | 33 | 4 | 33.33% |
| **C2 T2** | 11 | 11 | 56 | 7 | 63.64% |
| **C2 T3** | 12 | 15 | 40 | 7 | 46.67% |
| **C3 T1** | 1 | 7 | 30 | 3 | 42.86% |
| **C3 T2** | 2 | 14 | 33 | 8 | 57.14% |
| **C3 T3** | 3 | 14 | 22 | 7 | 50.0% |

Table 6: **Gene selection comparison of the timeBAE with a linear regression approach.** The second row consists of the collected gene selection information for all latent dimensions of the timeBAE and for all fitted linear models (latent dimensions were reorganized as described in Methods Section 5.7). Each alternate row represents the comparison between the linear model for the respective cluster and time point and the visually corresponding timeBAE latent dimension. timeBAE latent dimensions 7,8, and 9 were excluded. The percentages of gene overlap were calculated relative to the number of genes selected for the timeBAE latent dimensions.

#### S13 Simulation of stair-like scRNA-seq-like data

For the following analysis, we simulate a binary count matrix modeling 10 developmental stages using the simulation design from [4]. Specifically, a cell taking the value 1 for some gene indicates a gene expression level exceeding a certain threshold for that cell, whereas the value 0 indicates an expression level below that threshold. For $n = 1000$ cells and $p = 50$ genes, the data is modeled by i.i.d. random variables $(X_{i,j} \sim \text{Ber}(p_{i,j}))_{i=1,...,n,\ j=1,...,p}$, where $p_{i,j}$ denotes the probability of a value of $1$ for a gene $j$ in a cell $i$. In each developmental stage, the binarized expression levels of $100$ cells are generated, where $5$ genes are simulated as highly expressed, hence are characterizing for that cell stage, whereas the remaining genes are lowly expressed in that stage. Therefore, $p_{i,j}$ is set to 0.6 for highly expressed genes (HEGs) and to 0.1 for lowly expressed genes (LEGs). To model a smooth transition between two successive stages, HEGs of successive stages overlap, i.e., genes 1-5 characterize the first stage, genes 4-8 characterize the second stage, genes 7-11 the third stage, and so on, resulting in the stair-like structure shown in Supplementary Figure S12, Panel b. Details about the overlap patterns can be found in Supplementary Table 7. In total there are 32 HEGs and 18 lowly expressed genes across all cell stages which are added as non-informative genes to make the variable selection task more challenging and realistic.

| | Stage 1-2 | Stage 2-3 | Stage 3-4 | Stage 4-5 | Stage 5-6 |
|---|---|---|---|---|---|
| Overlap genes | 4, 5 | 7, 8 | 10, 11 | 13, 14 | 16, 17 |
| | Stage 6-7 | Stage 7-8 | Stage 8-9 | Stage 9-10 | |
| Overlap genes | 19, 20 | 22, 23 | 25, 26 | 28, 29 | |

Table 7: **Pairs of overlap genes between two successive cell stages**

#### S14 Simulation study to illustrate how the BAE identifies distinct cell groups and corresponding explanatory genes

To illustrate how the BAE works in a simple controlled scenario, we use simulated data. Specifically, our aim is to show that the BAE is capable of capturing distinct subgroups of cells in different dimensions together with characteristic sets of genes. We simulate scRNA-seq data with binarized counts indicating high vs. low expression of a gene as explained in Supplementary Section S13, similar to [4]. The simulation design mimics gene expression data where different cell types or stages are characterized by different sets of genes, with some overlap between the gene sets. We create a stair-like structure, shown in Supplementary Figure S12b, where 10 groups of cells are characterized by 5 highly expressed genes each, and there is some overlap in the gene sets (e.g., genes 1-5 characterize cell type 1, genes 4-8 characterize cell type 2, etc.). In addition to 32 cell type-characterizing genes, we simulate 18 non-informative "noise" genes. The simulated expression value for each cell is generated by sampling from a Bernoulli distribution with a higher success probability for a highly expressed gene, and a lower success probability for a lowly expressed gene (see Supplementary Section S13 for details). Data was standardized before model training and non-informative genes were re-scaled by a scaling factor of $\frac{2}{3}$.

We train a BAE with a 2-layer decoder and 10 latent dimensions. For evaluating the ability in extracting meaningful genes related to some groups of cells, we inspect the encoder weight matrix of the BAE during the training, since each nonzero weight represents a link between a gene and

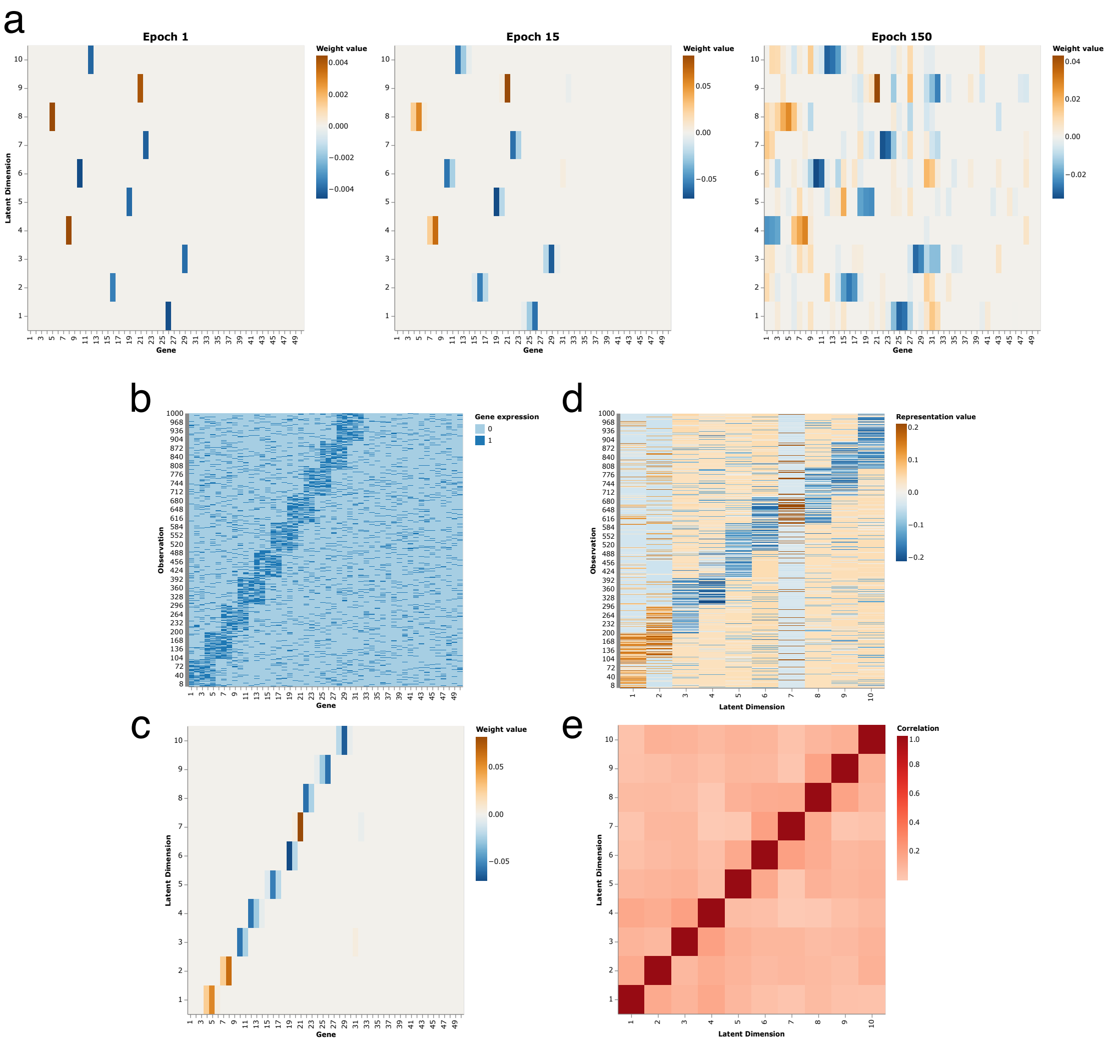


Figure S12: **BAE application on simulated scRNA-seq data.** **a** Heatmaps display the evolution of the BAE encoder weight matrix starting from a zero-initialization after training for 1, 15, 150 epochs (from left to right). **b** Heatmap of the simulated binary count matrix consisting of groups of cells in 10 different developmental stages. **c** We manually rearrange the latent dimensions of the encoder weight matrix after 15 epochs, so that learned patterns match the patterns in the count matrix for clarity. The heatmap displays the latent representation produced by the trained BAE after 15 epochs. **d** Heatmap shows the permuted encoder weight matrix of the BAE after training for 15 epochs. **e** Heatmap illustrates the absolute values of the sample Pearson correlation coefficients between the permuted latent dimensions of the BAE after 15 training epochs.

one latent dimension. The results are shown in Supplementary Figure S12. After the first training epoch, the BAE has selected one gene for each latent dimension, which serves as a guidance for further gene selection steps in the following epochs (Supplementary Figure S12a). During the next epochs, the model then learns to link additional genes to the distinct latent dimensions, characterizing the same group of cells as the already selected genes. When training proceeds for a larger number of epochs, at some point, the BAE starts updating additional weights corresponding to still underrepresented patterns. Stopping the training after e.g. 15 training epochs results in a sparse encoder weight matrix capturing the main characteristic patterns of the data (Supplementary Figure S12d). As overlap genes occur more frequently in the data than the genes exclusively abundant in one cell type, the model tends to preferentially select overlap gene pairs and each latent dimensions captures the two cell types associated via the overlap genes (Supplementary Figure S12c).

Training longer, also noise genes are selected, but with small weights (Supplementary Figure S12a). Supplementary Figure S12e shows the absolute values of Pearson correlation coefficients between the 10 latent dimensions of the BAE after training for 15 epochs. All off-diagonal entries are small, confirming that latent dimensions of the BAE capture distinct factors of variation. Since the BAE focused on patterns driven by overlap genes, two neighboring latent dimensions have a slightly higher correlation.

One main source of uncertainty in learning the sparse encoder weight matrix are the randomly sampled decoder parameters. Before training the model, the decoder parameters are randomly sampled from a probability distribution, e.g. a common strategy is to sample the parameters according to the normalized initialization strategy [5], where the decoder weights are sampled from a uniform distribution, while the biases are set to zero. We investigated stability of the BAE latent representation with respect to different initializations of the decoder parameters across several training runs. We found the encoder weight matrices to display very similar patterns and sparsity levels after 15 training epochs (see Supplementary Figure S13) and almost no non-informative genes are selected.

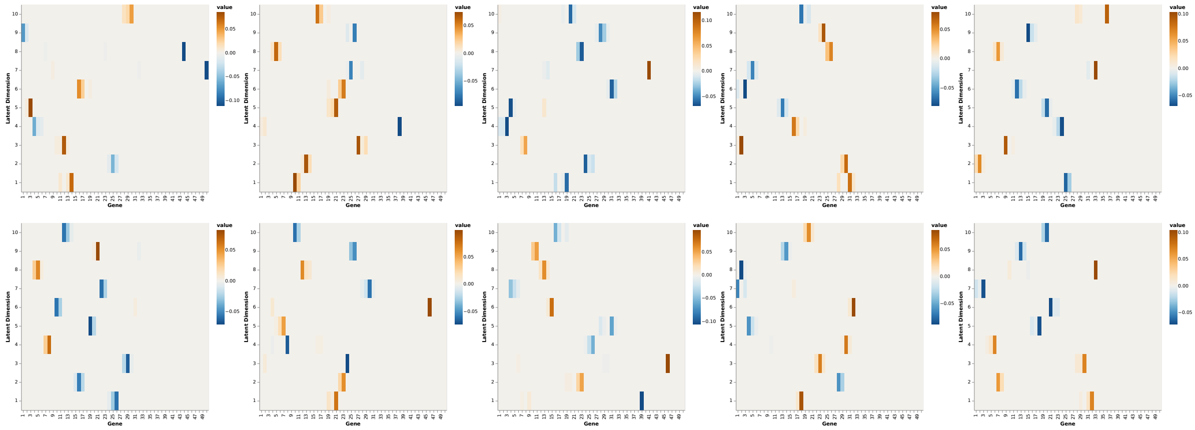

Figure S13: **BAE encoder weight matrices for different decoder parameter initializations.** Heatmaps represents the learned encoder weight matrices of BAEs after 15 training epochs with differently initialized decoder parameters. Training of the BAEs was performed using the same simulated dataset and hyperparameters.

#### S15 Model comparison of different autoencoder versions on simulated data

To demonstrate the effectiveness of the different components in our modeling approach for identifying a sparse set of explanatory genes and latent dimensions that encode distinct groups of cells, we compared our BAE approach to 1) a BAE without the disentanglement constraint, i.e., with unconstrained negative gradients as responses; 2) a standard unregularized autoencoder; 3) an autoencoder with an additional $L_1$ penalty term to enforce sparsity; 4) an autoencoder with an additional correlation penalty term to enforce uncorrelated latent dimensions; 5) a variational autoencoder (VAE) [6]. Detailed results are shown in Supplementary Figures S14-S19).

Specifically, in our comparison we consider a vanilla autoencoder (AE) using the mean squared error as the reconstruction loss

$$L_{\text{rec}}(\boldsymbol{x}, \widehat{\boldsymbol{x}}) := \frac{1}{p}\sum_{j=1}^{p}(x_j - \widehat{x}_j)^2,$$

$$L_{\text{AE}}(\boldsymbol{X}, \widehat{\boldsymbol{X}}) := \frac{1}{n}\sum_{i=1}^{n} L_{\text{rec}}(\boldsymbol{x}_i, \widehat{\boldsymbol{x}}_i),$$

an AE where we added a weighted $L_1$-penalty term for constraining the encoder parameters ($L_1$AE), such that the training objective which is to minimize becomes

$$L_{L_1\text{AE}}(\boldsymbol{X}, \widehat{\boldsymbol{X}}) := L_{\text{AE}}(\boldsymbol{X}, \widehat{\boldsymbol{X}}) + \alpha_{L_1\text{AE}} \cdot \sum_{j=1}^{p}\sum_{l=1}^{d}|\beta_{j,l}|,$$

and an AE where we added a weighted correlation penalty term utilizing the sample Pearson correlation coefficient $\widehat{\text{cor}}(\boldsymbol{x}, \boldsymbol{y})$ that penalizes the correlations between latent dimensions, such that the training objective which is to minimize becomes

$$L_{\text{corAE}}(\boldsymbol{X}, \widehat{\boldsymbol{X}}) := L_{\text{AE}}(\boldsymbol{X}, \widehat{\boldsymbol{X}}) + \alpha_{\text{corAE}} \cdot \sum_{k=1}^{d}\sum_{l=1}^{d}\widehat{cor}(f_{\text{enc}}(\boldsymbol{X};\ \boldsymbol{\psi})_k, f_{\text{enc}}(\boldsymbol{X};\ \boldsymbol{\psi})_l)^2,$$

where $\boldsymbol{\psi}$ denote the encoder parameters of the AE model.

In addition, we consider a VAE with a multivariate standard normal prior distribution for the latent space. The decoder learns the means of a multivariate normal distribution with a diagonal covariance matrix and fixed equal variance parameters $\sigma^2_{\text{dec}}$. The loss function for training the VAE is

$$\hat{\mu}_{ij},\ \hat{\sigma}^2_{ij} = f_{\text{enc}}(\boldsymbol{x}_i;\ \boldsymbol{\psi})_j,$$

$$L_{\text{VAE}}(\boldsymbol{X}, \widehat{\boldsymbol{X}}) := \frac{1}{2\sigma^2_{\text{dec}}}\sum_{i=1}^{n}\sum_{j=1}^{p}(x_{ij} - \widehat{x}_{ij})^2 - \frac{1}{2}\sum_{i=1}^{n}\sum_{j=1}^{p}(1 + \log\hat{\sigma}^2_{ij} - \hat{\mu}^2_{ij} - \hat{\sigma}^2_{ij}),$$

where $\boldsymbol{\psi}$ denote the encoder parameters of the VAE model and the second term on the right side of the equation corresponds to the Kullback–Leibler divergence.

For a fair comparison, we used the same architecture and training hyperparameters for the different (V)AE versions and for the BAE, except that the (V)AE versions are allowed to have a trainable bias vector in their single-layer encoders but no activation function. For training the different AE versions, we are using the Adam optimizer [7] and set the learning rate for the optimization to $0.01$. For training the VAE we use the AdamW optimizer [8], which incorporates weight decay regularization for the VAE encoder and decoder parameters. The weights for the different penalty terms were set to $\alpha_{L_1\text{AE}} = 0.1$, $\alpha_{L_1\text{AE}} = 0.075$ and $\alpha_{\text{VAE}} = 0.1$. The weight values were selected by empirically investigating the quality of the latent representation during the optimization process.

While there are no further restrictions made for the encoder of the AE, the $L_1$-penalty term in the $L_1$AE shrinks most of the weights towards zero during the optimization process, resulting in a sparse encoder weight matrix consisting of many values close to, but not exactly zero. Empirically, however, that happened only after many training epochs, such that the $L_1$AE is prone to overfitting, as opposed to the BAE, where starting from a zero initialization, meaningful variables are selected already during the first training epochs so that early stopping can be applied. The correlation penalty term incorporated in the loss function of the corAE enforces the model to learn a latent representation with uncorrelated latent dimensions.

Briefly, the BAE is the only model that identifies the cell group-characterizing genes already in the very first training epochs while maintaining uncorrelated latent dimensions that correspond to distinct groups of cells. The encoder weight matrix is sparse (Supplementary Figure S14, S15), due to the stagewise selection of the boosting component in the encoder, which starts from a zero-initialized weight matrix. In contrast, all the other models start training from a nonzero random initialization of the encoder weight matrix. As a result, a sparse (and hence interpretable) encoder weight matrix that represents meaningful patterns is learned much later in the training process or not at all (Supplementary Figures S16, S17, S18). The comparison with a standard BAE shows that the additional constraint on the selection criterion is essential for capturing distinct groups of cells in different latent dimensions and ensuring uncorrelated latent dimensions (Supplementary Figure S15).

Of the other models, only the corAE provides uncorrelated latent dimensions, which is not surprising given that its training objective explicitly penalizes correlation between latent dimensions. Yet, it does not learn a clear structure for the latent representation (Supplementary Figure S18). Similarly, the standard autoencoder does not learn structured latent representations (Supplementary Figure S16). While the $L_1$AE is able to find some cell stage characterizing patterns, which can be seen in the heatmaps of the encoder weight matrix and the latent representation after training for 1000 epochs (Supplementary Figure S17), most of the encoder weights are not truly zero, compromising interpretability. For the VAE with weight decay regularization, the encoder weight matrix is dense in every stage of the optimization process. However, the model seems to be more robust in capturing patterns during later training epochs (1000 and 25000 epochs), but the latent dimensions tend to be more correlated compared to the other models (Supplementary Figure S19).

In summary, while the correlation and $L_1$ penalties are effective in the respective separate models, the BAE achieves both characteristics at the same time without an additional penalty term. The uncorrelated latent dimensions are enforced only via modification of the selection criterion, which can be flexibly tailored to encode also other constraints, such as time structure, as shown in Section 2.4.

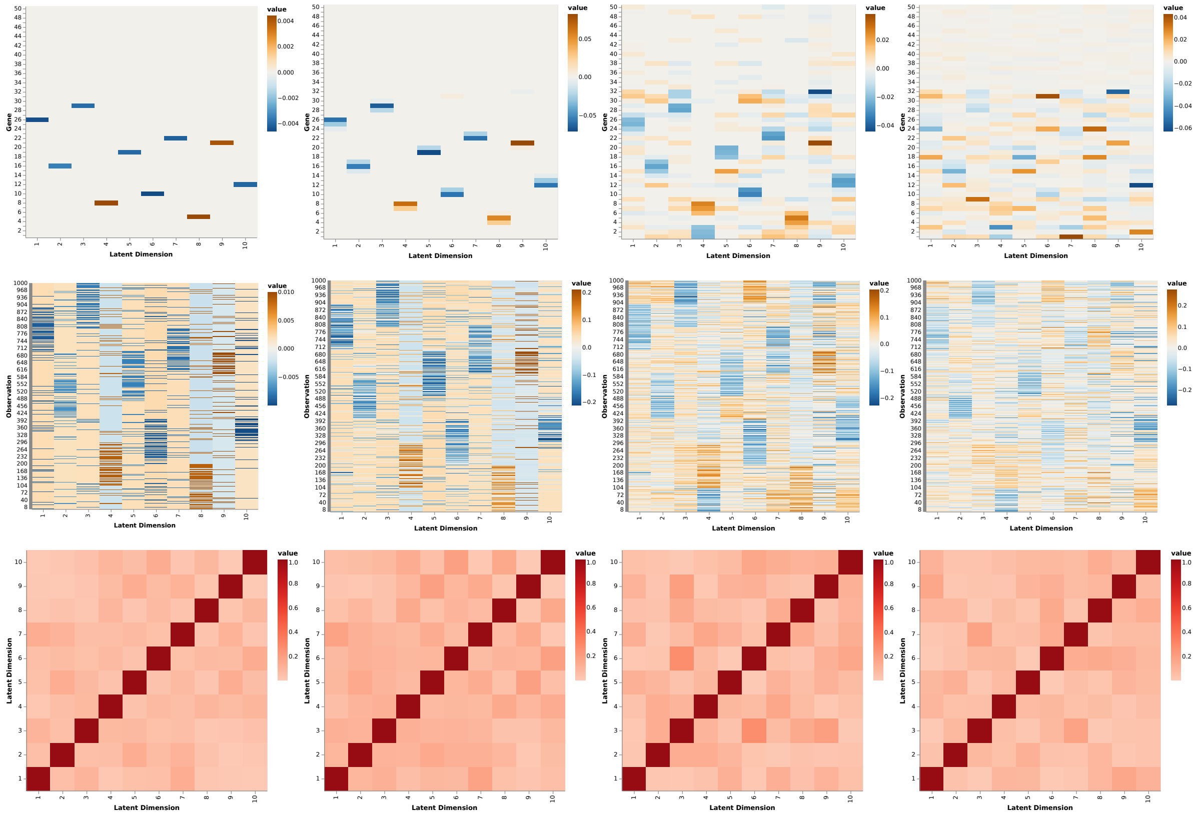


Figure S14: **BAE results on simulated scRNA-seq data during different training epochs.** Different rows display the heatmaps of the encoder weight matrix (top row), the latent representation (middle row) and the absolute sample Pearson correlation between the different latent dimensions (bottom row). Each column represents the results after training the model for a certain number of training epochs: 1, 15, 1000, 25000 (from left to right).

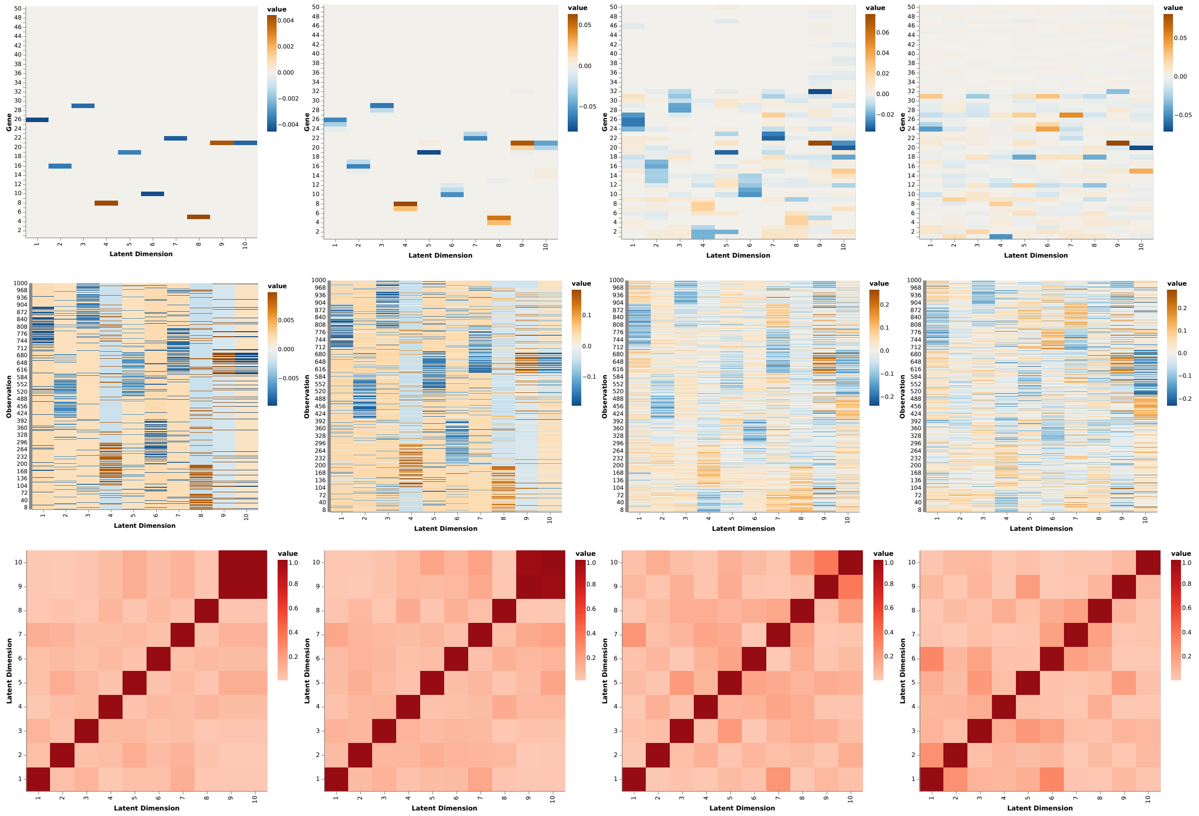

Figure S15: **BAE results without the disentanglement constraint on simulated scRNA-seq data during different training epochs.** Different rows display the heatmaps of the encoder weight matrix (top row), the latent representation (middle row) and the absolute sample Pearson correlation between the different latent dimensions (bottom row). Each column represents the results after training the model for a certain number of training epochs: 1, 15, 1000, 25000 (from left to right).

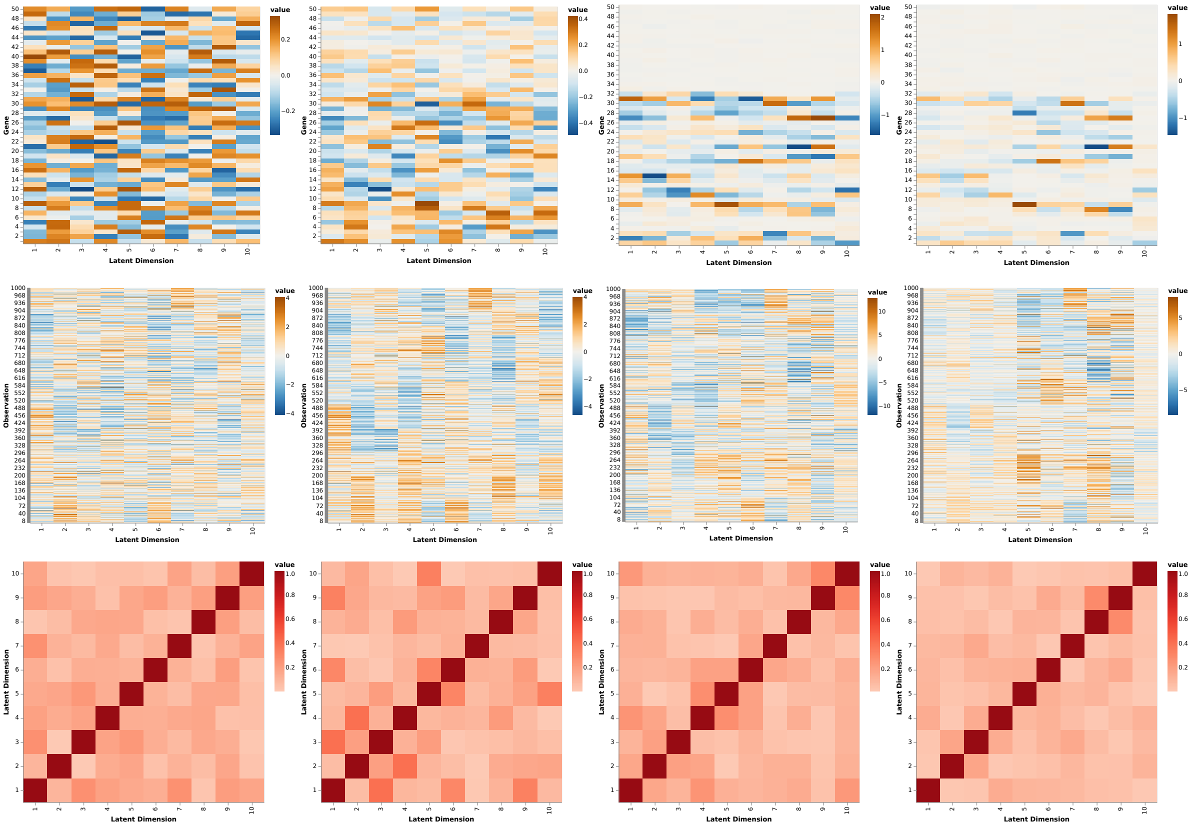


Figure S16: **AE results on simulated scRNA-seq data during different training epochs.** Different rows display the heatmaps of the encoder weight matrix (top row), the latent representation (middle row) and the absolute sample Pearson correlation between the different latent dimensions (bottom row). Each column represents the results after training the model for a certain number of training epochs: 1, 15, 1000, 25000 (from left to right).

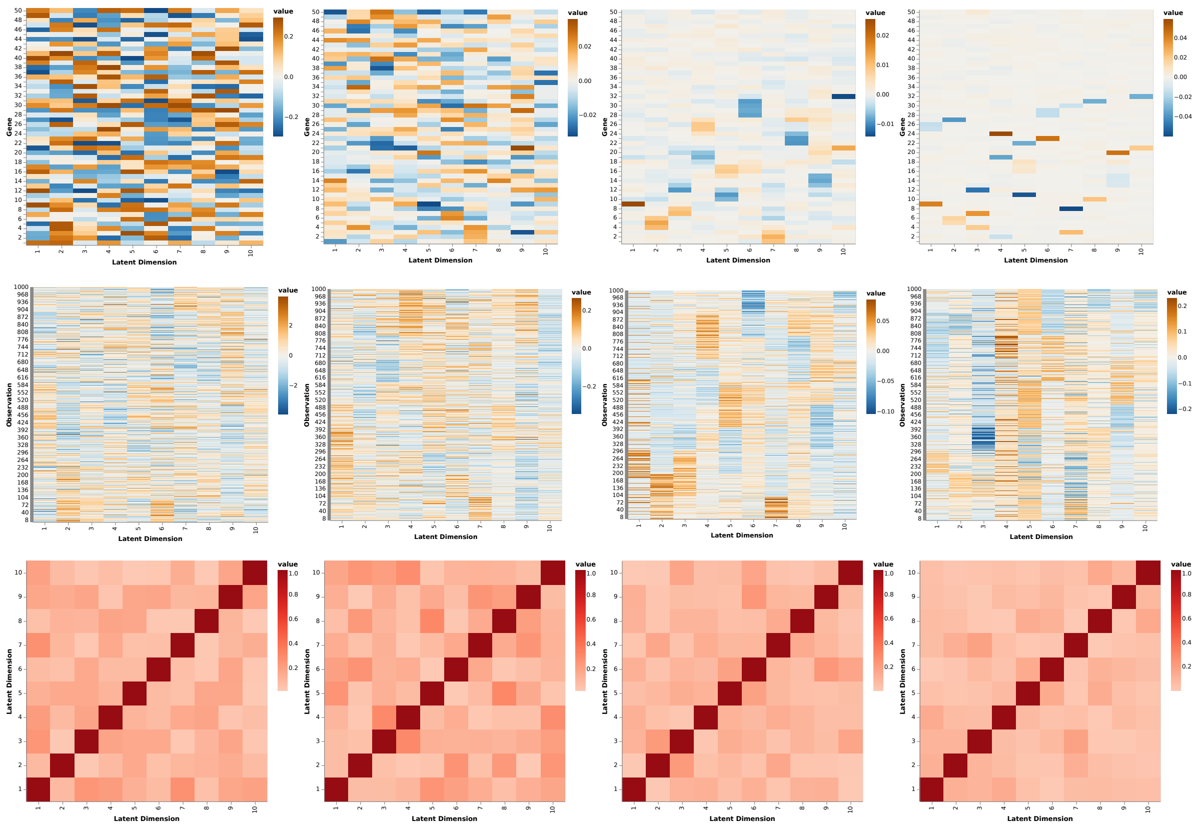

Figure S17: $L_1$**AE results on simulated scRNA-seq data during different training epochs.** Different rows display the heatmaps of the encoder weight matrix (top row), the latent representation (middle row) and the absolute sample Pearson correlation between the different latent dimensions (bottom row). Each column represents the results after training the model for a certain number of training epochs: 1, 15, 1000, 25000 (from left to right).

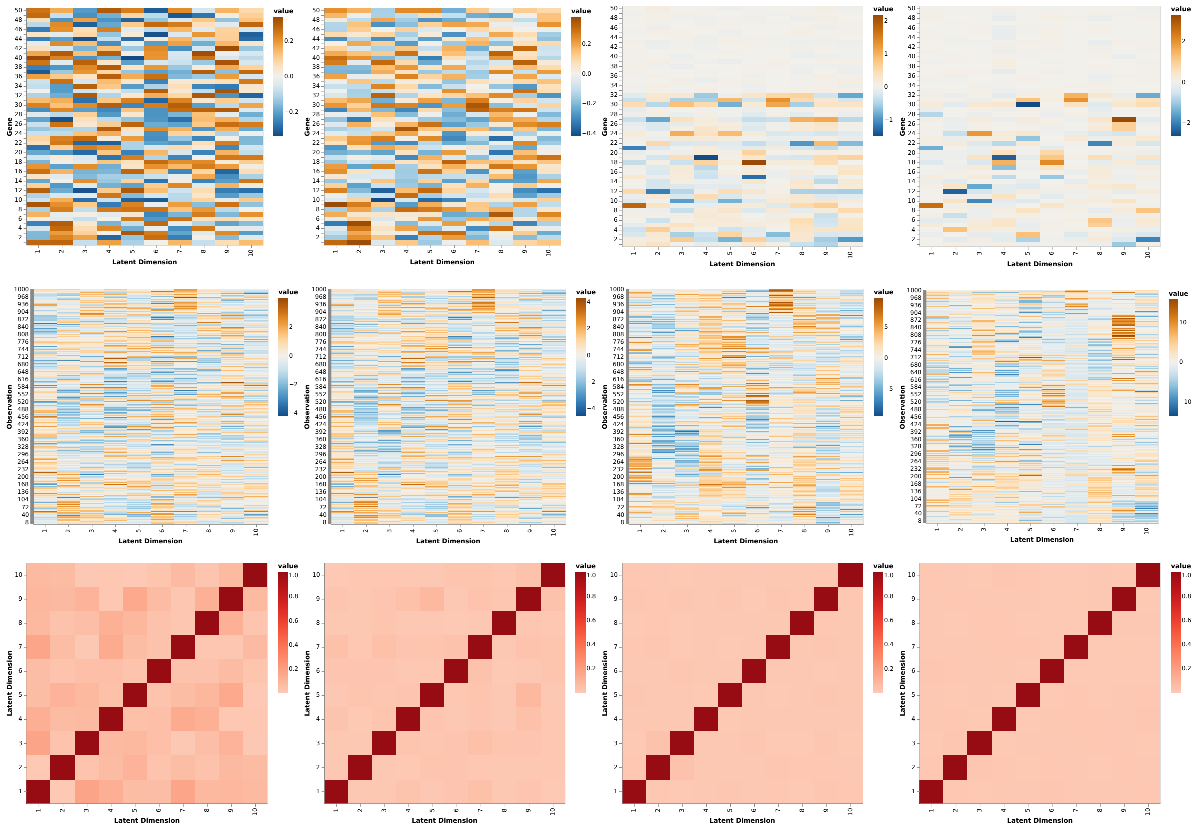

Figure S18: **corAE results on simulated scRNA-seq data during different training epochs.** Different rows display the heatmaps of the encoder weight matrix (top row), the latent representation (middle row) and the absolute sample Pearson correlation between the different latent dimensions (bottom row). Each column represents the results after training the model for a certain number of training epochs: 1, 15, 1000, 25000 (from left to right).

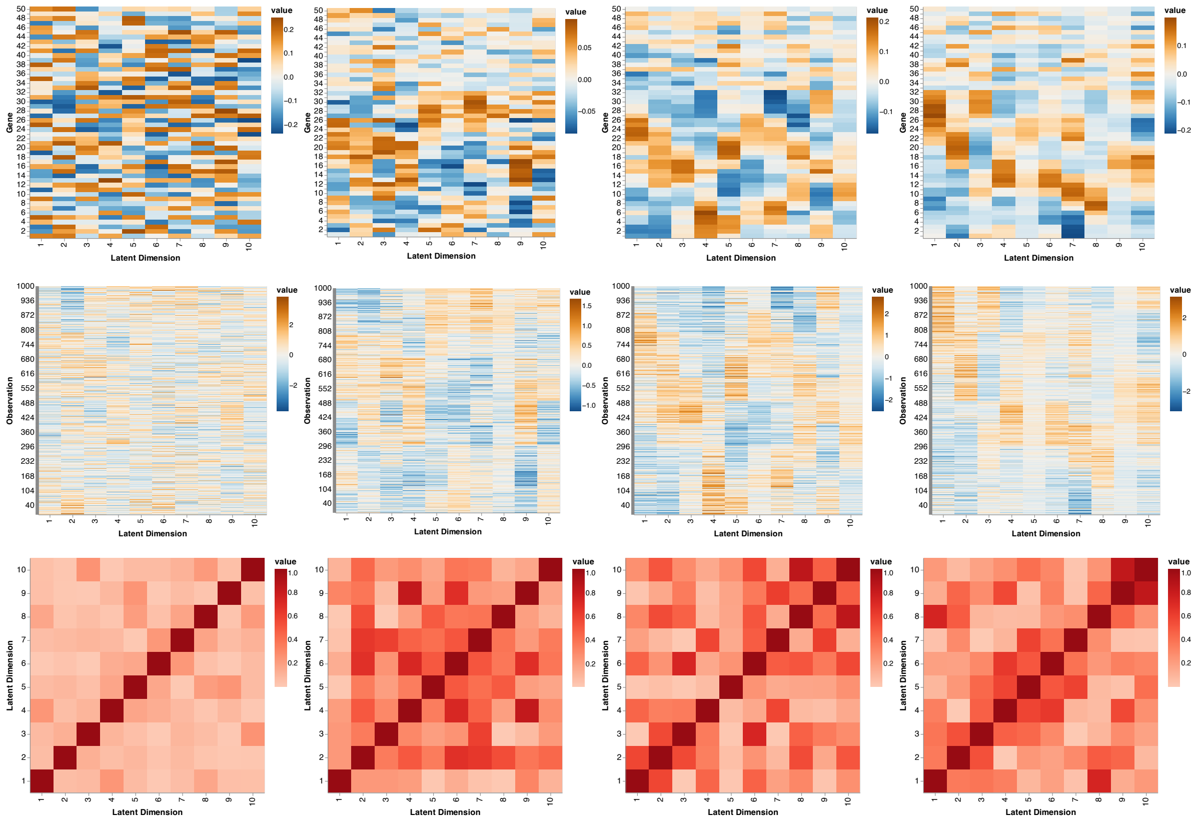


Figure S19: **VAE results on simulated scRNA-seq data during different training epochs.** Different rows display the heatmaps of the encoder weight matrix (top row), the latent representation (middle row) and the absolute sample Pearson correlation between the different latent dimensions (bottom row). Each column represents the results after training the model for a certain number of training epochs: 1, 15, 1000, 25000 (from left to right).

#### S16 Pseudocodes

**Algorithm 1:** comp$L_2$Boost algorithm

**Input:** Initial estimate $\widehat{\boldsymbol{\beta}}^{(0)} \in \mathbb{R}^p$, standardized observation matrix $\boldsymbol{X} \in \mathbb{R}^{n\times p}$, mean centered response vector $\boldsymbol{y} \in \mathbb{R}^n$, step size $\varepsilon \in (0,1)$, number of boosting iterations $M \in \mathbb{N}$

**Output:** Estimated solution $\boldsymbol{\beta}^*$

**for** $m = 1, \ldots, M$ **do**

$\widehat{\boldsymbol{\eta}}^{(m)} = \boldsymbol{X}\widehat{\boldsymbol{\beta}}^{(m-1)}$;

$\boldsymbol{r}^{(m)} = \boldsymbol{y} - \widehat{\boldsymbol{\eta}}^{(m)}$;

$\widehat{\boldsymbol{\gamma}}^{(m)} = \frac{1}{n-1} \cdot \boldsymbol{X}^\top \boldsymbol{r}^{(m)}$;

$j^* = \underset{j=1\ldots,p}{\arg\max}\, |\widehat{\gamma}_j^{(m)}|$;

$$\widehat{\boldsymbol{\beta}}^{(m)} = \begin{cases} \widehat{\beta}_j^{(m-1)} + \varepsilon \widehat{\gamma}_j^{(m-1)} & , j = j^* \\ \widehat{\beta}_j^{(m-1)} & \text{, else.} \end{cases};$$

**end**

$\boldsymbol{\beta}^* = \widehat{\boldsymbol{\beta}}^{(M)}$

**Algorithm 2:** Core BAE optimization algorithm

**Input:** Standardized observation matrix $\boldsymbol{X} \in \mathbb{R}^{n\times p}$, BAE with $d$ latent dimensions, zero-initialized transposed encoder weight matrix $\boldsymbol{B}$ and randomly initialized decoder parameters $\theta$, number of boosting iterations $M \in \mathbb{N}$, step size $\varepsilon \in (0,1)$ learning rate $\nu \in (0,1)$, batch size $m \leq n$

**Output:** Encoder weights $\boldsymbol{B}^*$

**while** *stopping criterion is not met* **do**

$\boldsymbol{Z} = \boldsymbol{X}\boldsymbol{B}$;

$\boldsymbol{g}^{(l)} = -\nabla_{\boldsymbol{z}^{(l)}} \frac{1}{n} \sum_{i=1}^n L_{\text{rec}}(\boldsymbol{x}_i, f_{\text{dec}}(\boldsymbol{z}_i;\ \theta)),\ l = 1, \ldots, d$;

**for** $l$ ***in*** $1, \ldots, d$ **do**

$\widehat{\boldsymbol{g}}^{(l)} = \text{standardize}(\boldsymbol{g}^{(l)})$;

$\boldsymbol{\beta}^{(l)} = \text{comp}L_2\text{Boost}(\boldsymbol{\beta}^{(l)}, \boldsymbol{X}, \widehat{\boldsymbol{g}}^{(l)};\ \varepsilon, \text{steps} = M)$;

**end**

$\boldsymbol{B} = (\boldsymbol{\beta}^{(1)}, \ldots, \boldsymbol{\beta}^{(d)})$;

$\boldsymbol{\theta} = \text{optimize}(\theta, m, \nu;\ \boldsymbol{B})$

**end**

$\boldsymbol{B}^* = \boldsymbol{B}$

**Algorithm 3:** Disentanglement constraint

**Input:** Number of latent dimension $l \in \{1, \dots, d\}$, standardized observation matrix $\boldsymbol{X}$, transposed encoder weight matrix $\boldsymbol{B} \in \mathbb{R}^{p \times d}$, standardized pseudo response vector $\widehat{\boldsymbol{g}}^{(l)} \in \mathbb{R}^n$

**Output:** Constraint pseudo response vector $\widehat{\boldsymbol{y}}^{(l)}$

$\mathcal{I} := \{l\} \cup \{k = 1, \dots, d \mid \beta^{(k)} = \mathbf{0}_{\mathbb{R}^p}\}$;

**if** $|\mathcal{I}| = d$ **then**

  $\widehat{\boldsymbol{y}}^{(l)} = \widehat{\boldsymbol{g}}^{(l)}$;

**end**

**else**

  $\boldsymbol{Z} = \boldsymbol{X}\boldsymbol{B}$;

  $\boldsymbol{\alpha} = ((\boldsymbol{Z}^{(-\mathcal{I})})^\top \boldsymbol{Z}^{(-\mathcal{I})})^{-1} (\boldsymbol{Z}^{(-\mathcal{I})})^\top \widehat{\boldsymbol{g}}^{(l)}$;

  $\boldsymbol{y}^{(l)} = \widehat{\boldsymbol{g}}^{(l)} - \boldsymbol{Z}^{(-\mathcal{I})} \boldsymbol{\alpha}$;

  $\widehat{\boldsymbol{y}}^{(l)} =$ standardize$(\boldsymbol{y}^{(l)})$;

**end**
